# Orchestration of SARS-CoV-2 Nsp4 and host-cell ESCRT proteins induces morphological changes of the endoplasmic reticulum

**DOI:** 10.1101/2024.12.08.627411

**Authors:** Allison Kifer, Franciso Pina, Nicholas Codallos, Anita Herman, Lauren Ziegler, Maho Niwa

## Abstract

Upon entry into the host cell, the non-structural proteins 3, 4, and 6 (Nsp3, Nsp 4, and Nsp6) of severe acute respiratory syndrome coronavirus 2 (SARS-CoV-2) facilitate the formation of double-membrane vesicles (DMVs) through extensive rearrangement of the host cell endoplasmic reticulum (ER) to replicate the viral genome and translate viral proteins. To dissect the functional roles of each Nsp and the molecular mechanisms underlying the ER changes, we exploited both yeast *S. cerevisiae* and human cell experimental systems. Our results demonstrate that Nsp4 alone is sufficient to induce ER structural changes. Nsp4 expression led to robust activation of both the unfolded protein response (UPR) and the ER surveillance (ERSU) cell cycle checkpoint, resulting in cortical ER inheritance block and septin ring mislocalization. Interestingly, these ER morphological changes occurred independently of the canonical UPR and ERSU components but were mediated by the endosomal sorting complex for transport (ESCRT) proteins Vps4 and Vps24 in yeast. Similarly, ER structural changes occurred in human cells upon Nsp4 expression, providing a basis for a minimal experimental system for testing the involvement of human ESCRT proteins and ultimately advancing our understanding of DMV formation.

## Introduction

The severe acute respiratory syndrome coronavirus 2, **SARS-CoV-2**, is a member of the beta- coronavirus family with a long positive-strand RNA genome, encoding 16 nonstructural proteins (Nsps) and four viral structural proteins^1,2,3^. Upon entry into host cells, SARS-CoV-2 rearranges the host cell endoplasmic reticulum (**ER**) membrane, a vital compartment in eukaryotic cells, to generate unique double-membrane vesicles (**DMVs**), enclosing the SARS-CoV-2 RNA genome^3–7^. The formation of DMVs is thought to protect the viral RNA genome from the host cell defense mechanisms, such as ribonucleases, and to spatially concentrate the components required for replication and translation of the viral genome to produce viral proteins. Studies have shown that expression of the ER-localized proteins of either SARS-CoV-2, or the related flavivirus or Dengue virus in human tissue culture cells causes ER morphological changes, ultimately leading to the assembly of DMVs or DMV-like structures^8–10^. While recent studies have begun to unveil functional roles of these ER-localized Nsp proteins, the molecular mechanisms of DMV formation remain largely elusive.

The ER is one of the biggest compartments in eukaryotic cells and has several unique features. Structurally, it is continuous with the nuclear envelope (NE) and extends throughout the cytoplasm to form a reticular structure^11–16^. Functionally, it performs a variety of pivotal tasks, including generating almost all secretory pathway proteins, synthesizing the early stages of most lipids, and serving as a storage organelle for cellular calcium. In addition to its diverse roles, the ER’s functional capacities must adapt in response to environmental and developmental cues. This is achieved in part by a signal transduction pathway, the unfolded protein response (UPR), which ensures that the size, shape, and functional capacities of the ER align with the cell’s demands^17–19^. Exceeding the functional capacity of the ER and its various functions is considered a contributor of canonical “ER stress”, and the UPR upregulates the transcription of genes encoding proteins involved in ER protein folding and modifying components. Permanently misfolded proteins are ultimately degraded through ER-associated degradation (ERAD)^20–25^. Independent of the UPR and ERAD, ER functional homeostasis is also regulated during cell division in yeast cells. We discovered a cell cycle checkpoint that ensures the inheritance of sufficient functional ER in the two daughter cells in dividing yeast, *S. cerevisiae,* called the ER surveillance (**ERSU**) pathway^25–31^. Upon ER stress induction, the entry of the ER into the daughter cell is blocked and the septin ring becomes mislocalized from the bud neck to the bud scar (the site of a previous cell division), leading to a halt in the cell cycle^27^. Once the ER functions are replenished through the UPR pathway, cells are released from this blockage and re-enter the cell cycle.

The yeast ER exhibits a relatively simpler structure^32,33^, with the perinuclear ER (pnER) consisting of an outer nuclear membrane (ONM) that is contiguous with the inner nuclear membrane (INM), although the nuclear pore complex (NPC) separates the INM from the pnER (or ONM). Additionally, yeast ER contains cortical ER (cER), juxtaposed with the plasma membrane, and the tubular ER connects the cER with the pnER. Although the precise details of the morphological changes and functions remain elusive, the ER undergoes changes depending upon the functional needs^30,34^, influencing ER homeostasis. Furthermore, the ER cannot be generated *de novo*. Thus, the functions, overall levels, and size of the ER must be properly regulated during the cell cycle by the ERSU pathway, which operates independently from the UPR^25.^

The ability of the ER lipid bilayer to adjust its structure and functional capacities is exploited by pathogens such as SARS-CoV-2 and other RNA viruses to generate DMVs for the replication of viral particles within the host cell. The significance of DMV formation is underscored by the fact that an inability to form DMVs prevents viral proliferation in closely related RNA viruses such as Middle East respiratory syndrome and SARS-CoV^7,35^. Despite our knowledge that the SARS-CoV-2 proteins Nsp3, Nsp4, and Nsp6 are involved in DMV generation, relatively little has been elucidated about the molecular mechanisms by which these viral proteins interact with host cell components to achieve this. To dissect these molecular events and their individual roles, we expressed SARS-CoV-2 Nsp4 under a regulatable promoter in yeast, leveraging its simpler ER architecture, which provides immense advantages in detecting subtle morphological changes. Additionally, the ease of genetic manipulation in yeast makes it an ideal system for identifying host genes that contribute to ER morphological changes in coordination with Nsp4. Utilizing this minimal system as a proof of principle, we found that Nsp4-induced ER changes are mediated by proteins of the endosomal sorting complex required for transport (ESCRT), which are known for their roles in a wide range of membrane processes, including cytokinesis, ER micro-autophagy, and NE formation^36^. Our results revealed that the expression of SARS-CoV-2 Nsp4 causes distinct effects on the functions, morphology, and inheritance of ER during the cell cycle. We also tested whether our findings in yeast are conserved in human cells. Together, we anticipate that our results will lay the foundation for a genome-wide screen to identify host proteins responsible for the ER morphological changes that ultimately lead to DMV formation.

## Results

Three SARS-CoV-2 Nsp proteins predicted to be localized to the ER, Nsp3, Nsp4, and Nsp6, have been implicated in DMV formation. To dissect the functional roles of individual Nsp proteins and the mechanisms of DMV formation, we established an experimental system in yeast where SARS-CoV-2 Nsp4 is expressed under a galactose (Gal)-regulatable promoter. We hypothesized that, in comparison to the architecture of the ER in mammalian cells, the simpler ER morphology of yeast cells would likely facilitate robust observations of the ER structural changes induced by Nsp4. While we did not expect expression of a single Nsp protein to generate the full scope of DMV formation, we reasoned that certain features of the ER structural changes, such as ER membrane bending, scission, establishment of new ER links, or ER ligation, might lead to DMV formation. Dissecting the functional roles of Nsp4, together with Nsp3 and Nsp6, will help unravel the molecular mechanisms of SARS-CoV-2 DMV formation as well as the concurrent ER structural changes.

To this end, we visualized both the cER and pnER of wild-type (WT) yeast upon integration of a well-characterized optical reporter, Pho88-GFP, an ER-membrane protein fused with green fluorescent protein (GFP), at the genomic PHO88 locus (we refer to these cells as WT Pho88-GFP cells) (**Figure 1A**). Without Nsp4 expression, Pho88-GFP depicted the pnER surrounding the nucleus and the cER below the cortex of the plasma membrane with a relatively simple continuous ER morphology, as observed previously^26^. We next transformed a yeast 2μ plasmid carrying the SARS-CoV-2 Nsp4-Flag gene placed under a Gal-inducible promoter into WT Pho88-GFP cells. We examined the impact of Nsp4 expression on the ER morphology by shifting the growth media that contain 2% raffinose (-Gal) to that with 2% Gal (+Gal) for 3 hr at 30°C (**Figure 1A-B**). Consistent with previous reports^31^, we observed a relatively uniform distribution of Pho88-GFP for both the pnER and the cER without Nsp4 expression in the raffinose-containing media **(Figure 1A-B, Fig *S1A*).** However, upon inducing Nsp4 expression in the Gal-containing media, confirmed by Western blot (**Figure *S1B***), the overall uniform and continuous morphologies of both the cER and pnER were significantly altered. Specifically, certain portions of both the cER and pnER showed foci or accumulation of Pho88-GFP (**Figure 1B**, +Gal, marked by white arrowheads), scattered between the discontinuous cER and pnER. We rarely observed such ER morphology with GFP foci when cells were grown in the raffinose-containing media, indicating that Nsp4 expression caused significant Pho88-GFP distribution changes. As Nsp4 is an ER-membrane protein, we tested whether overexpression of Nsp4 induced the UPR pathway. For this experiment, we used a well-established UPR reporter, UPRE-GFP, where GFP is placed under an UPR transcription enhancer element (UPRE). Upon shifting to the Gal-containing media, we observed GFP expression, revealing that Nsp4 overexpression resulted in activation of the UPR (**Figure *S1C***).

**Figure 1:**
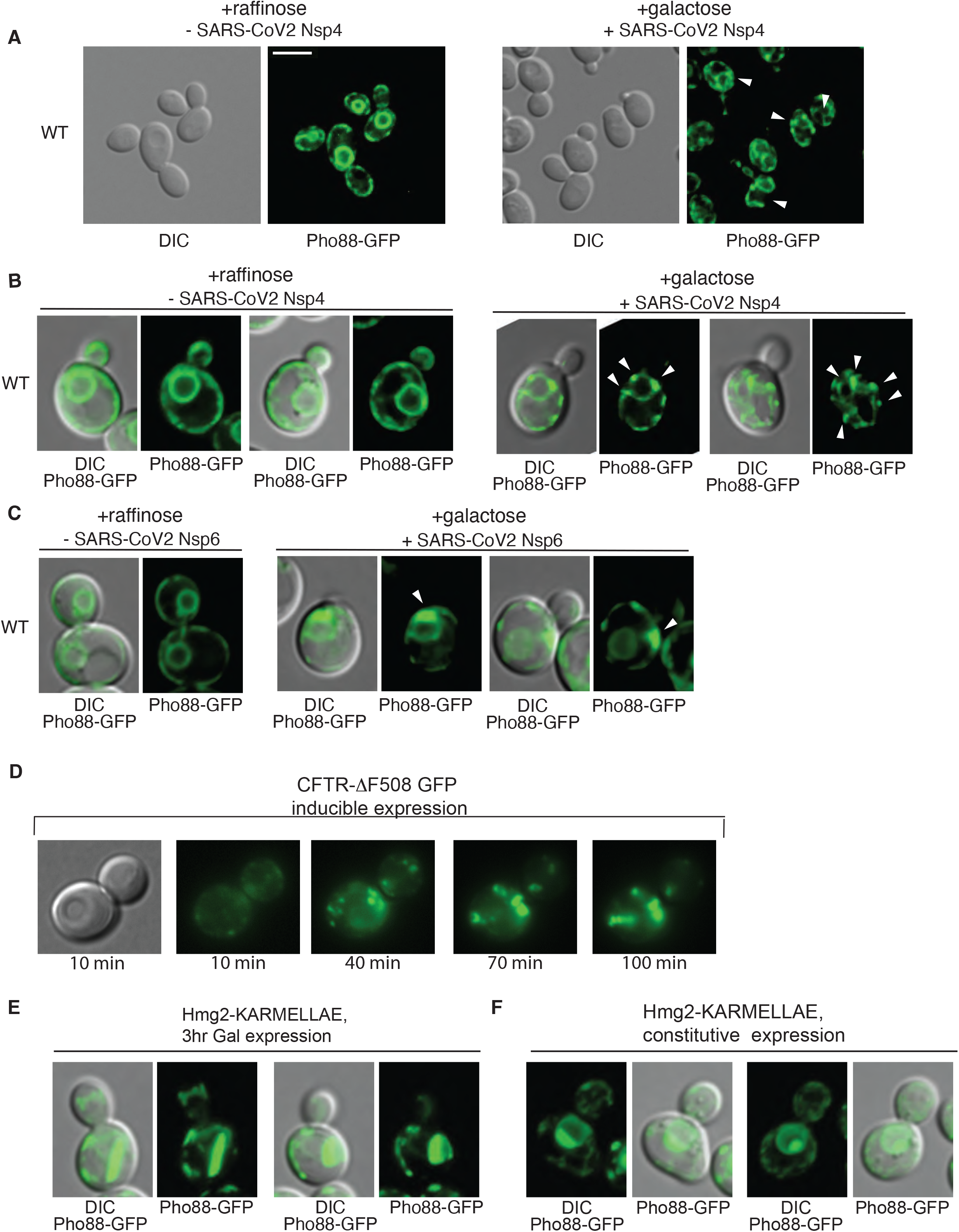
SARS-CoV-2 Nsp4 expression leads to the yeast ER morphological change. **(A)** The ER of WT yeast cells carrying an ER-membrane optical reporter, Pho88-GFP, was visualized in the absence (left) or presence (right) of SARS-CoV-2 Nsp4 expression. Upon transformation of a plasmid carrying SARS-CoV-2 Nsp4 placed under a Gal-inducible promoter, cell growth was shifted from raffinose (left) to Gal (right) containing media for 3 hr before visualization of the ER. For both conditions, differential interference contrast (DIC) and fluorescence microscopy (Pho88-GFP, green) images are shown. White arrowheads point to some of the Pho88-GFP-accumulated foci. Scale bar, 5 μm. **(B)** Close-up images of the ER of cells initially grown in raffinose (left), and then shifted to Gal-containing media (right) White arrowheads highlight Pho88-GFP aggregation/foci. **(C)** Close-up images of the ER, visualized by the ER Pho88-GFP reporter, in the absence (left) and presence (right) of SARS-CoV-2 Nsp6**. (D)** CFTR-ΔF508-GFP aggregates increased over time upon shifting to Gal-containing media. **(E)&(F)** Karmellae were visualized by the Pho88-GFP reporter upon transient expression of Hmg2 by shifting to Gal-containing media for 3 hr **(E)** or constitutive expression upon growth in glucose media **(F).** In both cases **(E)** and **(F)**, the overall morphological changes by Hmg2 appear similar. However, dramatic changes were diminished with constitutive expression of Hmg2, indicating that cells presumably adapted towards Hmg2 expression.

The distribution of Pho88-GFP within the ER was also altered upon expression of another SARS-CoV-2 ER transmembrane protein, Nsp6 (**Figure 1C**). However, the alterations differed from those caused by Nsp4, suggesting that each ER-membrane SARS-CoV-2 Nsp might impose distinct effects on the ER. It should be noted, however, that Nsp6 was constitutively expressed, and thus, some of the observed changes may be due to differences in expression methods, whether induced (Nsp4) or constitutively expressed (Nsp6). When Nsp4 expression was continued overnight, however, the Pho88-GFP distribution remained similar to that observed with a 3-hr induction of Nsp4. Importantly, the morphological changes of the Pho88-GFP distribution were not unique to the Pho88- GFP ER optical reporter, but rather represent the overall ER morphological changes, based on the use of the ER tracker that visualizes the ER membrane (data not shown).

Over the years, several morphological changes of the ER membrane have been reported. To this end, we tested whether the ER morphological changes caused by Nsp4 resemble those reported with overexpression of human cystic fibrosis transmembrane conductance protein, CFTR-FD508, which is commonly mutated in humans. Overexpression of CFTR-FD508 showed distinct puncta, consistent with our previous observations (**Figure 1D**)^30^. Additionally, overexpression of an ER-membrane protein, Hmg1-CoA reductase, led to accumulation of Hmg1-GFP adjacent to the nucleus (**Figure 1E**). Previous electron microscopy (EM) studies have reported a unique ER structure called *Karmellae*^37^, with stacks of the ER membrane partially enclosing the nucleus. Using Hmg1-GFP, we observed that the ER morphological changes differed from those observed with Nsp4 expression. Notably, although the specific features were distinct, the overall extent of morphological changes was relatively comparable with those observed with Nsp4-Flag (**Figure 1E**). Interestingly, the ER morphological changes induced by the regulatable overexpression of Hmg1-GFP (i.e., *Karmellae)* were more pronounced than those seen with the constitutive expression of Hmg1-GFP, suggesting potential adaptation in ER structural changes (**Figure 1E-F**). Taken together, the expression of either Nsp4 or Nsp6 alone in WT host cells can lead to ER morphological changes that are distinct in specific features from *Karmellae* but similar in the overall extent of the changes.

### Nsp4 expression induces cER inheritance block

In addition to the UPR, we previously reported that ER stress independently induces the ERSU cell cycle checkpoint pathway in yeast (**Figure *S1D***)^26^. Under normal growth conditions, a daughter cell starts to emerge from the mother cell, and the tubular ER originating from the pnER points to the incipient bud tip or the bud tip and enters the emerging daughter cell (**Figure *S1E*)**. To facilitate our analyses, we categorized cells into three classes depending on the cell cycle stage. Class I cells are defined by a bud index (the ratio of daughter cell diameter to mother cell diameter) of less than 0.3, as described previously^26^. As daughter cells grow, class II cells see an increase in the bud index to between 0.3 and 1.0. The initial tubular ER that enters the bud extends towards the bud tip. Upon attachment to the polarisome at the bud tip, the initial tubular ER growth shifts laterally, generating class II cells with the cER spreading under the cortex of the daughter cell. Finally, in class III cells, as daughter cells continue to grow, both the pnER and the nucleus enter the daughter cell (with a bud index >0.3–1), followed by nuclear division. Previously, with canonical ER stress (e.g., tunicamycin [Tm] treatment-induced ER stress), we reported that the mechanisms of cER entry into the daughter cell resulted in two distinct outcomes for ER inheritance block, depending on when the ER stress is recognized (**Figure *S1E-F***): (1) If a cell detects ER stress early in the cell cycle, prior to the initial ER attachment to the polarisome at the bud tip, stable ER entry into the daughter cell is blocked. This is followed by septin ring mislocalization, ultimately leading to cytokinesis block (**Figure *S1F;* ER inheritance block 1**). In contrast, (2) if cells encounter ER stress after the initial ER attachment to the polarisome at the bud tip, the cER does not go back into the mother cell but continues to spread under the cell cortex, with both the pnER and the nucleus entering the daughter cell without septin ring mislocalization, leading to cell division upon cytokinesis (**Figure *S1F;* ER inheritance block 2**). In those divided daughter cells, the cell cycle proceeds to produce class I cells, where cER inheritance is blocked in the second round of cell division (**Figure *S1F;* ER inheritance block 2**).

Based on these findings, we anticipated that the activation of ER inheritance block in response to overexpression of viral proteins such as Nsp4 would behave similarly to ER inheritance in response to canonical ER-stressed cells treated with Tm, a glycosylation inhibitor agent. However, upon shifting to Gal-containing media, we found that Nsp4 expression induced ER inheritance block in all three classes of cells (**Figure 2A-B**). The observed differences in the ER inheritance block with Nsp4 expression were unlikely to be due to differences in growth media, because we observed cER inheritance block with Tm at similar levels as cells grown in either dextrose- or raffinose-containing media (**Figures *S2A-D***). Thus, the ER inheritance block in all cell classes with Nsp4 contrasts sharply with that induced by ER stress from Tm treatment. Indeed, with Tm treatment, we found that cER inheritance into class I and II daughter cells was blocked effectively, resulting in ∼50% of class I and class II cells lacking cER, while the cER inheritance block was least effective in class III cells (**Figure 2C**). This is in agreement with our previous results showing that once the ER is stably inherited, it is unable to be “un-inherited”^26^ (**Figure *S1F***).

**Figure 2:**
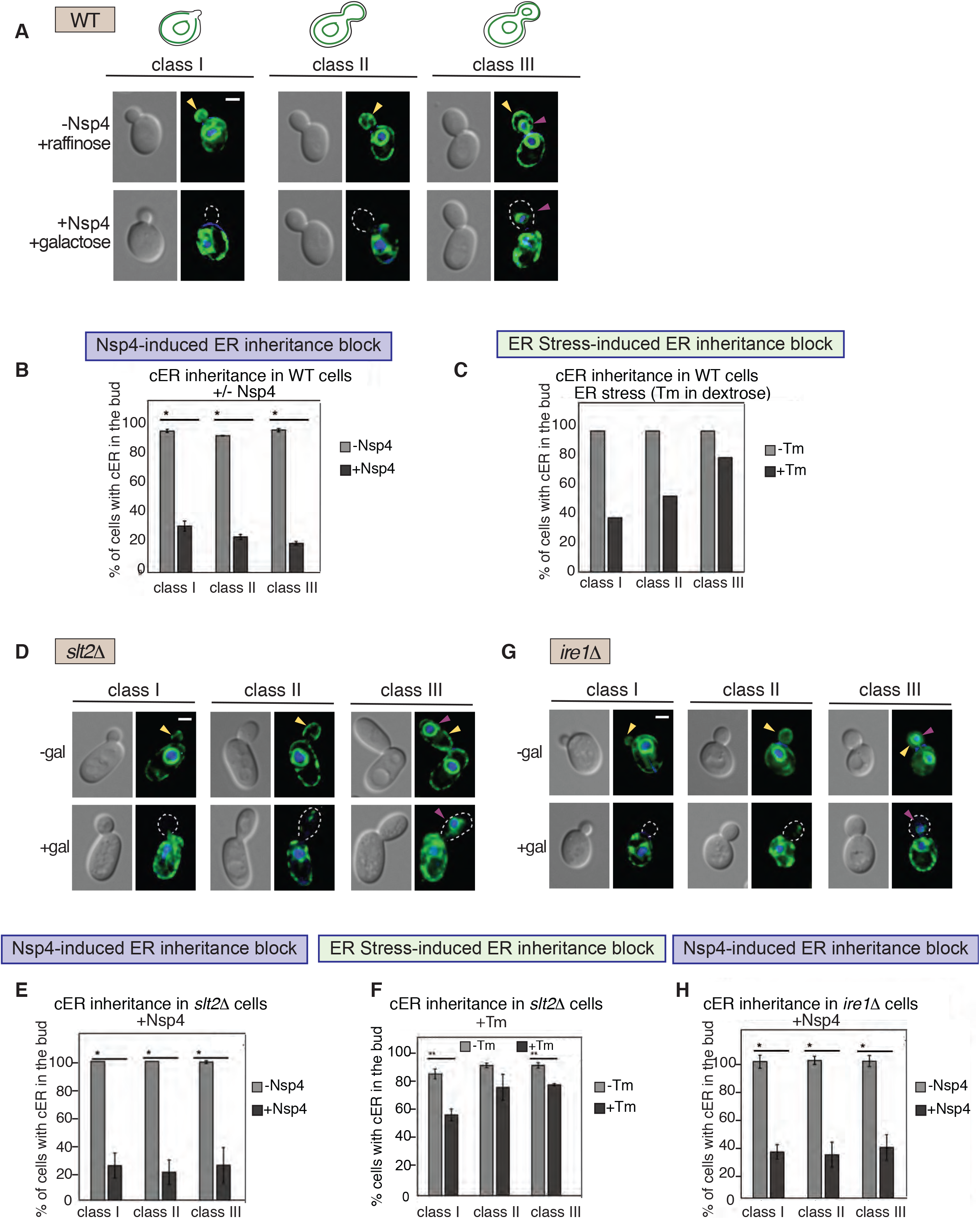
Nsp4 induces ER inheritance block via an unprecedented mechanism **(A)** ER inheritance block in class I, II, and III WT cells expressing an ER optical reporter, Pho88-GFP, and Gal-induced SARS-CoV-2 Nsp4 expression. Class I represents cells with small buds (bud index: <0.33), class II represents cells without perinuclear ER (pnER) (bud index: 0.33– 1), and class III represents cells with pnER in the daughter cell (bud index: 0.33–1), as described previously ^26^. Yellow arrowheads point to the cER and the purple arrowheads point to the pnER in the daughter cells. White dotted lines show the position of the daughter cell surface, demarked by tracing the DIC image. (**B**) Quantification of (A). Graph shows percentage of class I, II, and III WT daughter cells that inherited the cER, with (dark gray) or without (light gray) Nsp4 expression. (**C**) cER inheritance of WT cells treated with tunicamycin (Tm), a well-established ER stress- inducing drug, for 2 hr. Quantification of cER inheritance in WT cells with (dark gray) or without (light gray) Tm. (**D**) Images of ER inheritance in *slt2Δ* cells grown in raffinose (-Gal, -Nsp4) or Gal (+Gal, +Nsp4). (**E**) Quantification of (D). Graph shows percentage of class I, II, and III *slt2Δ* daughter cells that inherited the cER, with (dark gray) or without (light gray) Nsp4 expression. (**F**) Quantification of cER inheritance in *slt2Δ* cells with (dark gray) or without (light gray) Tm. (**G**) Images of ER inheritance in *ire1Δ* cells grown in raffinose (-Gal, -Nsp4) or Gal-containing media (+Gal, +Nsp4). (**H**) Quantification of (G). Graph shows percentage of class I, II, and III *ire1Δ* daughter cells that inherited the cER, with (dark gray) or without (light gray) Nsp4 expression. All graphs show the mean ± standard deviation (SD). ** indicates *p* < 0.05 and * indicates *p* < 0.01.

In addition to ER inheritance block, our previous work showed that proteotoxic ER stress results in septin ring mislocalization, moving from the normal bud neck to the bud scar or the cytokinetic remnant (CRM)^26,27^ (**Figure *S1D***). As the septin ring plays a critical role in cytokinesis, its mislocalization ultimately leads to cytokinesis block in yeast. Upon shifting the growth media from raffinose to Gal to express Nsp4 in WT cells expressing a GFP-tagged Shs1 septin subunit (Shs1-GFP) integrated at the Shs1 genomic locus^27^, we found that ∼25% of the Shs1-GFP was mislocalized from the bud neck to the CRM (visualized by staining with calcofluor white ([CFW]) (**Figure *S2E-I***). Tm treatment of WT cells in raffinose induced ∼45% mislocalization of Shs1-GFP **(Figure *S2E vs. S2F, and S2I*)**. Finally, even when Tm and Nsp4 expression were both induced, the extent of the Shs1-GFP mislocalization remained at ∼40% **(Figure *S2H-I*).** Interestingly, while most of the septin rings displayed mislocalization to the CRMs, we also observed separate and smaller fragments of Shs1-GFP foci localized outside of the bud neck or the CRMs **(Figure *S2G-H*,** see white arrowheads). Taken together, these results show that Nsp4 expression results in the activation of the ERSU molecular events, including both cER inheritance block and septin mislocalization. Nsp4-induced cER inheritance block occurs differently from the canonical ERSU event^26^. Even in class III cells, where the cER is anchored at the bud tip and/or the cER extends under the cortex of the daughter cells, the ER inheritance block occurs in all cell classes. These results suggest that the mechanisms underlying the cER inheritance block involve a different mechanism.

### Nsp4-induced ER inheritance block occurs independently of the ERSU pathway

Previously, we reported that the ERSU pathway occurs independently of the UPR pathway, involving a unique set of components such as SLT2 and reticulons (i.e., RTN1 and YOP1)^26,29^. To further investigate whether the Nsp4-induced cER inheritance block requires the ERSU component SLT2, we induced Nsp4 expression in *slt2D* cells upon switching to Gal-containing media (**Figure *S1A* and Figure 2D-E**). We found that expressing Nsp4 in *slt2Δ* cells grown in Gal-containing media caused cER inheritance block at a similar level as Nsp4 expressed in WT cells (**Figure 2B vs. 2E**). Strikingly, the ER inheritance block occurred in all three classes of *slt2D* cells, including class III cells with large daughter cells. These results revealed that Nsp4-induced ER inheritance block occurs effectively in class III cells in either WT or *slt2D* cells (**Class III cells in Figure 2B vs. 2E**). This contrasted with the behaviors of class III cells in response to Tm-induced ER stress (**Class III cells in Figure 2C vs. 2F)**. Taken together, these results validated the involvement of SLT2 in the canonical ERSU pathway as previously reported (specifically, for class I and II cells)^26^, while revealing that Nsp4-induced ER inheritance block occur differently without SLT2 involvement.

As the UPRE-GFP reporter was induced upon Nsp4 expression (**Figure *S1C***), we also tested the potential involvement of IRE1 in Nsp4-induced ER inheritance block. We found that Nsp4 expression in *ire1D* cells effectively caused cER inheritance block (**Figure 2G-H and Figure *S1B***). Together, these results revealed that a canonical ERSU component, SLT2, and a UPR component, IRE1, are not involved in the Nsp4-induced ER inheritance block. This highlights a unique feature of Nsp4-induced ER stress and ER inheritance block, distinct from previously described ER stress, despite the induction of the UPRE-GFP reporter (**Figure *S1C***). Taken together, our results uncovered that (1) Nsp4 expression induces cER inheritance block regardless of the cell cycle stage, even when cells encounter ER stress after the cER is stably inherited, and that (2) this block is mediated by a unique mechanism distinct from the canonical ERSU and UPR pathways.

### Nsp4-induced ER morphological changes are mediated by the ESCRT proteins Vps4 and Vps24

In addition to the ERSU molecular events, we also found that Nsp4 expression caused ER morphology changes even in *slt2D* and *ire1D* cells to a similar extent as WT cells upon switching to the Gal-containing media (**Figure 3A-C**). The cER and pnER morphologies were deformed in all three classes of cells upon Nsp4 expression in either *slt2D* (**Figure 2D and 3B**) or *ire1D* cells (**Figure 2G and 3C**), revealing that neither SLT2 nor IRE1 are involved in Nsp4-induced ER morphological changes. This result prompted us to conduct further investigation based on our current knowledge of ER functional and structural homeostasis. Towards this goal, we conducted a pilot screen by testing genes previously reported or suspected to be involved in the morphological changes of the ER membrane, such as in ERAD (e.g., Ubx2)^38,39^, the ER membrane lipid biosynthetic pathway (e.g., Mga2)^40^. In addition, we tested genes involved in ESCRT, including Vps4, Vps24 Vps23, and Snf7, as recent studies have shown that some of these genes play critical functions in lipid bilayer changes during ER microautophagy ^41–43^.

**Figure 3:**
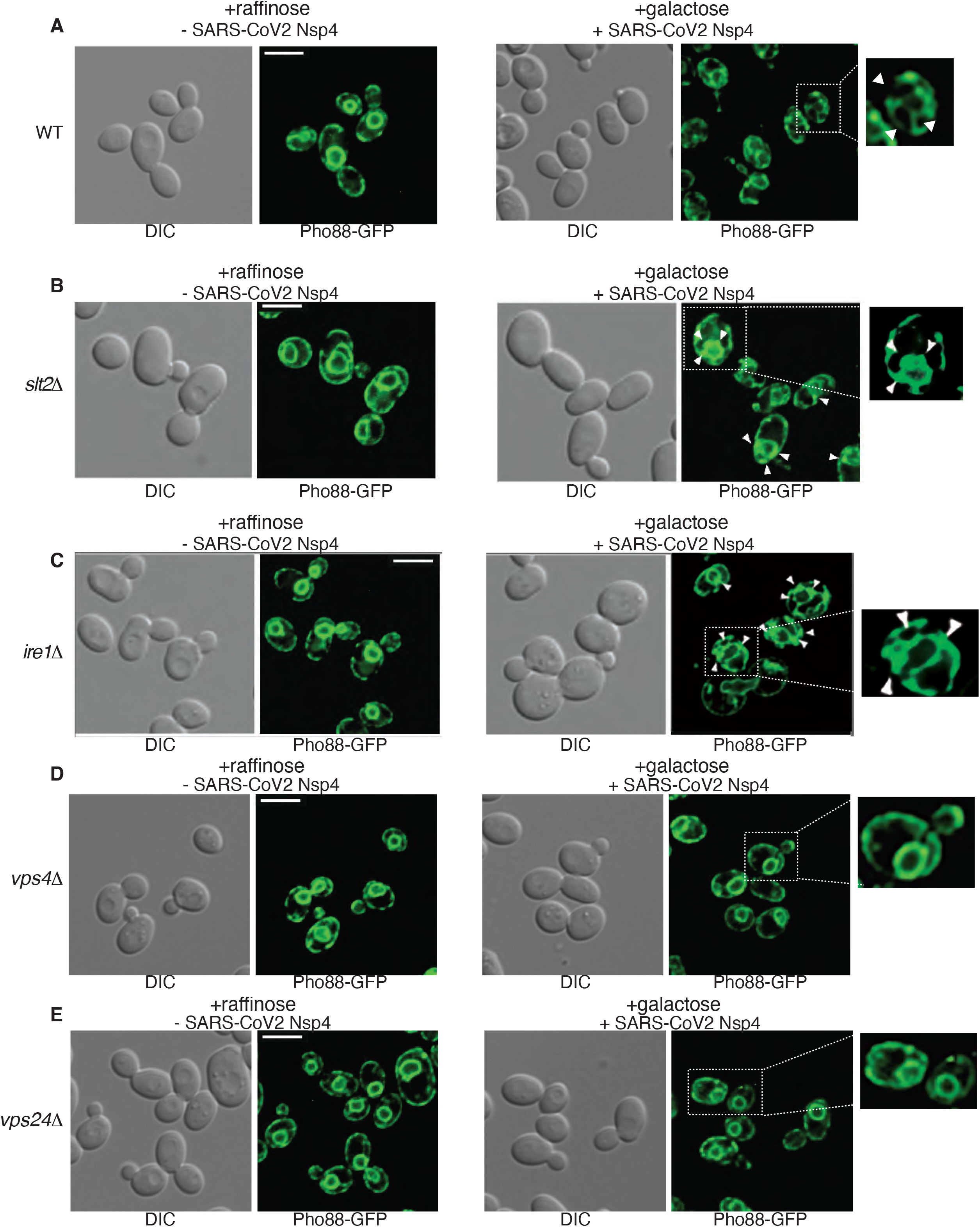
The ESCRT genes VPS4 and Vps24 play roles in the Nsp4-induced ER morphological changes. **(A)** Representative images of WT cells expressing Pho88-GFP in the absence (+raffinose) or presence (+Gal) of SARS-CoV-2 Nsp4 expression. Similarly, the ER was visualized in (**B**) *slt2Δ*, (**C**) *ire1Δ*, (**D**) *vps4Δ*, and (**E**) *vps24Δ* cells in the absence (+raffinose) and presence (+Gal) of SARS-CoV-2 Nsp4 expression. White arrowheads point to ER-structural changes and/or Pho88- GFP-accumulated foci. In *vps4Δ* or *vps24Δ* cells, Nsp4 expression did not cause significant morphological alterations of the ER and appeared similar to the ER in cells grown in media without Nsp4 expression. All scale bars, 5 μm.

Thus, as in our experiments in *slt2D* or *ire1D* cells (**Figure 3A-C**), we tested the ER morphology in *vps4D*, *vps24D*, *vps23D*, and *snf7D* cells upon transformation of Nsp4 under a regulatable promoter (**Figure 3D-E and *S3A-C***). If any of these genes were involved, we anticipated that the loss of ER morphological changes would occur even with Nsp4 expression. Upon shifting to Gal-containing media, we found that Nsp4 expression did not result in significant changes to the overall ER morphology in *vps4Δ* or, to a lesser extent, *vps24D* cells, unlike in WT cells (**Figure 3A vs. 3D-E**). Specifically, the distribution of both the cER and pnER in the *vps4Δ* cell became uniform with little foci, similar to the ER morphology found in WT cells in raffinose media without Nsp4 expression (**Figure 3A vs. 3D-E**). Note that Gal induction resulted in Nsp4 expression in both *vps4Δ* and *vps24Δ* cells (**Figure *S1A***); thus, the lack of ER foci was not due to the absence of Nsp4. Furthermore, not all ESCRT gene knockout cells showed recovery of the ER morphology. In contrast to *vps4Δ* or *vps24Δ* cells, the ER structural changes in *vps23Δ* and *snf7Δ* cells occurred similarly to those in WT cells in Gal-containing media (**Figure *S3A-C*).**

In addition to the overall recovery of ER morphology, we noticed that some *vps4Δ,* and to a lesser extent *vps24Δ,* daughter cells inherited the cER (**Figure 3D-E**). Thus, we tested whether the recovery of Nsp4-induced ER morphological changes in *vps4Δ* or *vps24Δ* cells also restored the ER inheritance block phenotype upon expression of Nsp4 (**Figures 4A-D**). ER inheritance occurred normally in *vps4Δ* and *vps24Δ* cells prior to Nsp4 expression. Upon Nsp4 expression, induced by shifting to Gal-containing media, cER inheritance was diminished to ∼60% of class I, ∼50% of class II, and ∼35% of class III *vps4Δ* cells **(Figure 4A-B**). These values indicate a partial recovery of the ER inheritance block compared to the Nsp4-induced WT cells, where cER inheritance was diminished to 30% in class I, 25% in class II, and 20% in class III cells (**Figure 4B**, WT). However, the cER inheritance block remained least effective in class I *vps4Δ* cells compared to class III cells, contrasting with the observation that the cER inheritance block was most pronounced in class I rather than class III cells in Tm-induced WT cells (**Figure 2C**). As reported previously, the ERSU-induced ER inheritance block occurred effectively before cER inheritance is committed upon attachment to the polarisomes, i.e., class I cells (**Figures *S1E-F***)^31^. Interestingly, the cER inheritance block in *vps23Δ* and *shs7Δ* cells upon Nsp4-GFP expression was similar to that of *vps4Δ or vps24Δ* cells (**Figure 4E-H**). Taken together, these results revealed that the ER morphological restoration and partial recovery of the cER inheritance block occur in the absence of Vps4, demonstrating a role of Vps4 in ER morphology and a partial role in the cER inheritance block induced by Nsp4.

**Figure 4:**
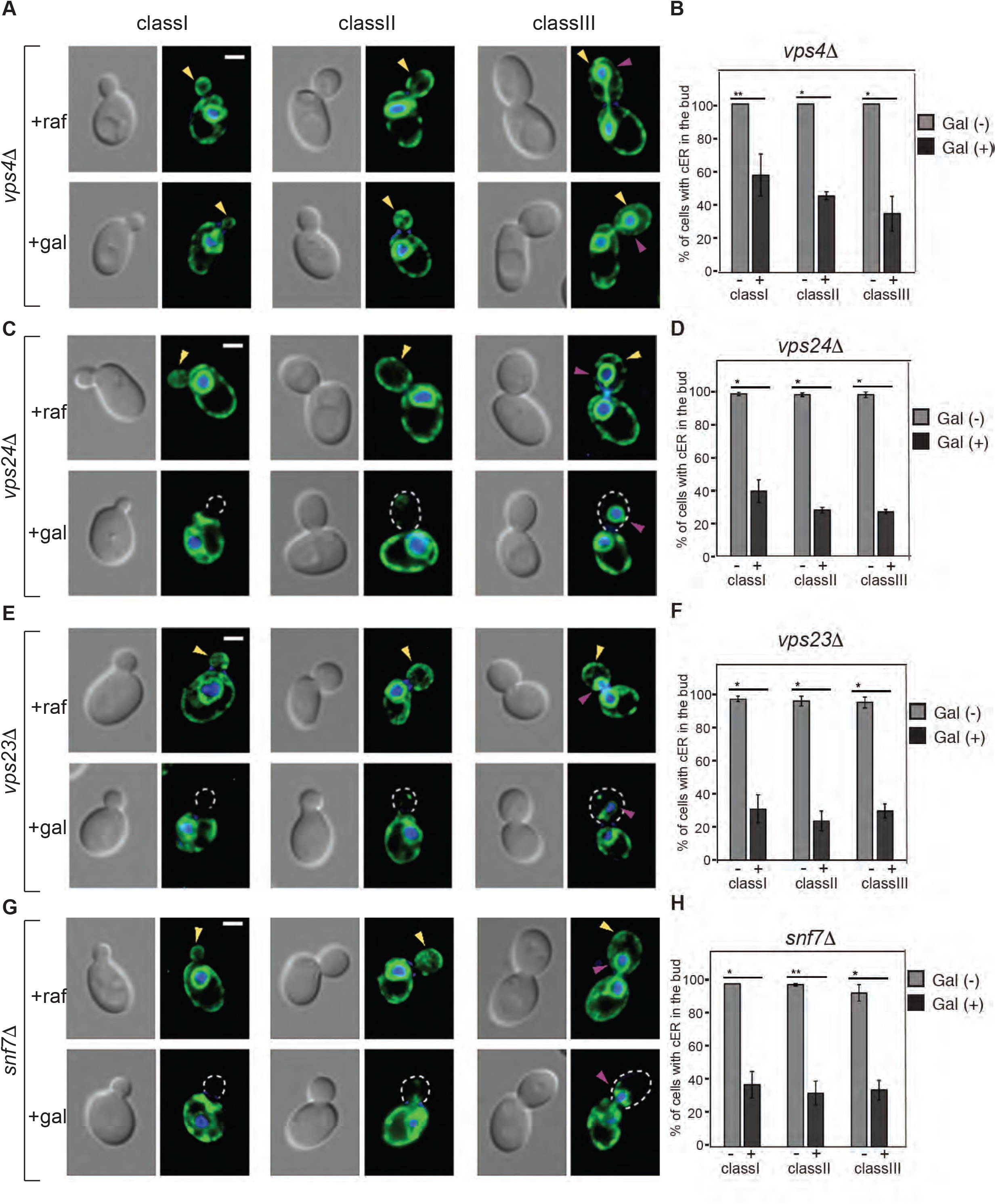
Nsp4-induced cER inheritance block was partially restored in *vps4Δ* cells. (**A**) cER inheritance block is induced by Nsp4 expression in *vps4Δ* class I, II, and III cells. Yellow arrowheads point to daughter cells with cER and purple arrowheads point to the pnER. (**B**) Quantification of cER inheritance block upon Nsp4 expression in *vps4Δ* class I, II, and III cells in comparison to those of WT cells. (**C**) cER inheritance block induced by Nsp4 expression in *vps24Δ* class I, II, and III cells and (**D**) quantification of the cER inheritance block in *vps24Δ* cells. (**E**)&(**G**) cER inheritance block in Nsp4-induced *vps23Δ* (**E**) and *snf7Δ* (**G**) cells. (**F**)&(**H**) Quantification of the cER inheritance block in (**E**) and (**G**). ** indicates *p* < 0.05 and * indicates *p* < 0.01.

### The impact of Nsp4 expression on the mammalian ER morphological structure

The robustness of our yeast experimental system using Nsp4, a SARS-CoV-2 gene, highlights the value of a minimal experimental system for investigating the roles of ER-localized SARS-CoV2 proteins individually. While SARS-CoV2 viral entry into human lung cells via the ACE2 receptor^44^ ultimately leads to DMV formation, we focused on investigating Nsp4-dependent ER morphological changes. To this end, we transfected a plasmid expressing Nsp4-RFP under a strong cytomegalovirus promoter into U2OS cells and monitored the ER morphology using a well-characterized human Sec61b-GFP ER optical reporter stably integrated into the genome of HeLa cells (**Figure 5A**) or U2OS cells (**Figure 5E**). We found that Nsp4 expression caused the formation of small circular foci-like or aggregates-like structures scattered within the ER network (**Figures 5A-B**, +Nsp4, indicated by white arrowheads and **Figure 5D**; quantitation of circular structures). Nsp4-RFP expression was confirmed by RFP levels in cells transfected with Nsp4-RFP expression upon performing fluorescence-activated cell sorting (FACS) analyses (**Figure 5C**). A similar structure was also observed with SARS-CoV-2 Nsp6 expression (**Figure 5B**, +Nsp6). Furthermore, the formation of these small foci-like structures induced by Nsp4 was independent of the ER optical reporter; we observed similar ER foci with a different ER reporter, Rtn4A-GFP, as well as ER tracker staining (**Figures *S4A-C*** *for Rtn4A and **Figures S4C and D** for ER tracker*). Thus, the morphological alterations observed with Nsp4-RFP expression are independent of the ER optical reporter utilized. Furthermore, longer Nsp4-RFP expression increased the levels of small foci-like or aggregates-like ER structures as expected (data not shown).

**Figure 5:**
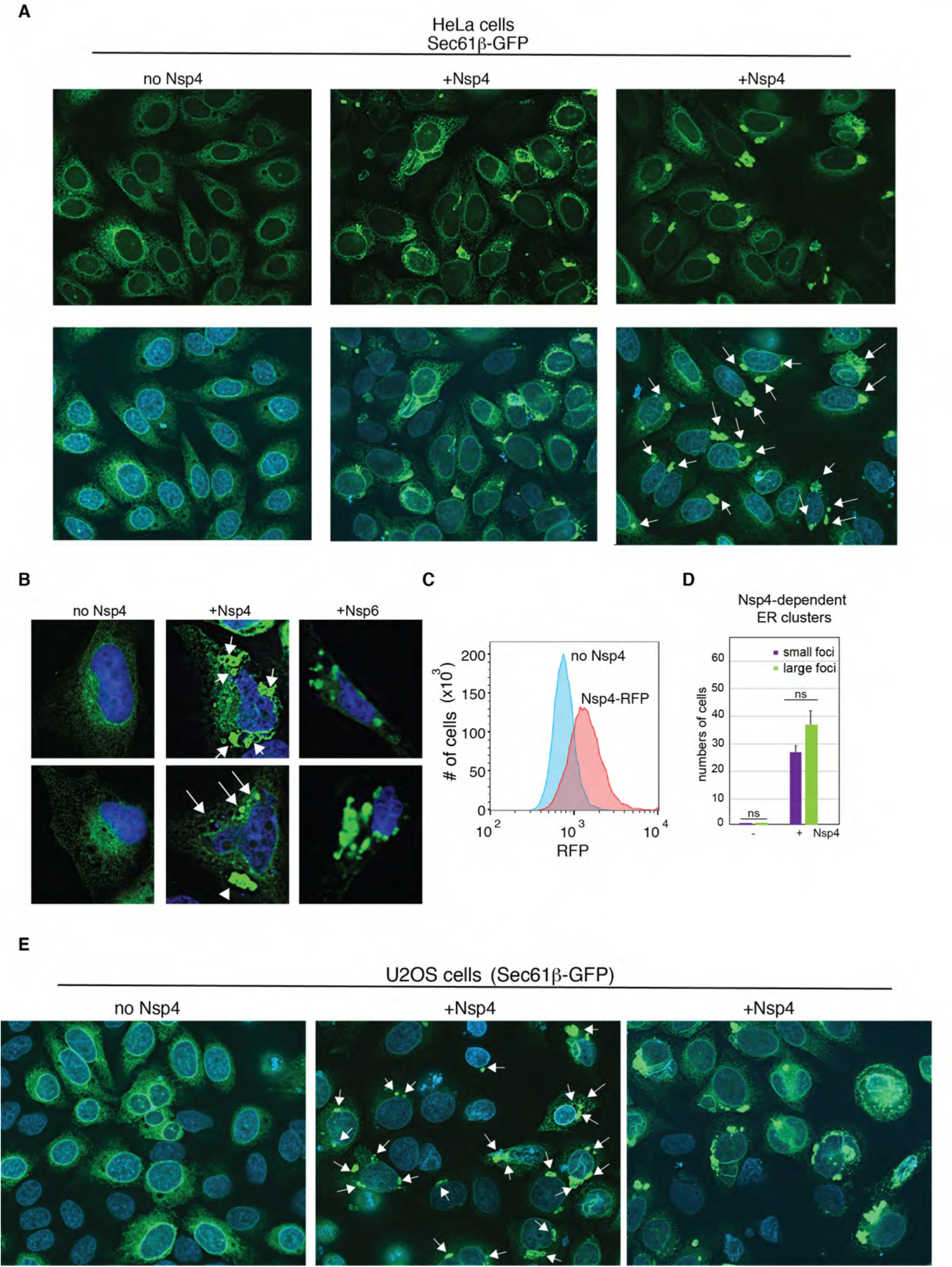
SARS-CoV-2 Nsp4 and Nsp6-induced ER morphological changes occur in human cells. **(A)** Representative field images of HeLa cells carrying a stably integrated Sec61β-GFP ER reporter with (+Nsp4) or without (no Nsp4) Nsp4 expression. Upper panels show Sec61β-GFP (green) and lower panels show merged images with DAPI-stained DNA. Nsp4 expression led to some of the ER morphological changes including generation of ER cluster/foci are highlighted with white arrows. (**B**) Close-up images of the ER morphologies using the Sec61b-GFP ER reporter upon constitutive expression of Nsp4 (+Nsp4), Nsp6 (+Nsp6), or no expression of Nsp4 or Nsp6 (no Nsp4) in HeLa cells. (**C**) Expression of Nsp4 in cells was analyzed by measuring Nsp4-GFP with Fluorescence-activated cell sorter (FACS) in comparisons to fluorescence levels in cells without Nsp4 expression. (**D**) Numbers of cells with Nsp4-induced ER circular-like structure changes (small foci (purple) and large foci (light green)) were quantitated. (**E)** Representative field images of U2OS cells carrying a stably integrated mEmerald-Sec61β ER reporter with (+Nsp4) or without (no Nsp4) Nsp4 expression and nuclear DNA is stained with DAPI. Some of the ER morphological changes, including ER aggregates/cluster are shown with white arrows.

To further characterize the impact of Nsp4 expression, we investigated whether Nsp4 expression impacted the morphologies of other secretory compartments or distributions of certain ER landmarks such as Sec31^45^, COPII vesicle formation sites (**Figure *S4A***), and ER–Golgi intermediate compartments (by staining with anti-ERGIC antibody) (**Figure *S4C***). We found that the distributions of Golgi-localized proteins, such as GPP130^46^ and GM130^47^, were altered although at lesser extent. In some cases, these proteins appeared protruded into the nucleoplasmic area or in the nucleus (**Figure *S4B and S4D&E***). Thus, Nsp4 expression caused architectural alterations of the early secretory pathway organelles in addition to the ER. Additionally, staining with 4′,6 diamidino-2-phenylindole (DAPI) revealed that the nuclear DNA exhibited deformed structures from the characteristic oval nuclear shapes; in some cases, we found DAPI-positive foci/fragments within and outside of the cell, suggesting that Nsp4 induced the damage or fragmentation of chromosomal DNA. Taken together, these observations revealed that expression of Nsp4 alone can effectively induce changes in the morphologies of the ER, and the early secretory pathway compartment in human cells.

## Discussion

The formation of DMVs is an effective strategy employed by positive-strand RNA viruses, such as those in the Flaviviridae family, which includes hepatitis C virus^48^, MERS, and SARS-CoV^3,49^, as a protective measure for their RNA genomes against host cell defense mechanisms like ribonucleases. Furthermore, DMVs concentrate host components necessary for efficient viral replication and translation, thereby promoting effective viral proliferation. However, despite recent intriguing findings^6,8^, the molecular mechanisms and identities of host cell components involved in DMV formation remain largely elusive, aside from SARS-CoV-2 proteins Nsp3, Nsp4, and Nsp6, which localize to the ER and play key roles in DMV formation. To better understand Nsp4-dependent ER structural changes and uncover the molecular mechanisms underlying DMV formation, we took a bottom-up approach by investigating the effects of SARS-CoV-2 Nsp4 expression on ER functions and morphology. Our research demonstrated that Nsp4 expression alone was sufficient to induce structural changes in the ER, including UPR induction and hallmark ERSU events such as ER inheritance block and septin mislocalization. Furthermore, we found that these ER changes occurred independently of both Ire1 and Slt2, canonical yeast cell components of the UPR and ERSU, respectively. Instead, this process is mediated by ESCRT proteins, Vps4 and, to a lesser extent, Vps24. Thus, our results suggest that SARS-CoV-2 Nsp4 collaborates with specific ESCRT proteins in host cells to alter ER morphology, and demonstrated the value of the minimum experimental system. The detailed mechanisms by which these ESCRT proteins function in Nsp4-induced ER structure in yeast remain to be investigated. Additionally, studying Nsp3 and Nsp6 in conjunction with Nsp4 will provide further insights into how ER-localized SARS-CoV-2 proteins alter ER architecture and form DMVs in human cells, potentially leading to new drug development strategies.

Interestingly, using SARS-CoV-2 protein Nsp4 revealed previously unrecognized differences in ERSU phenotypes. The impact of ER morphological changes caused by Nsp4 differed significantly from those caused by canonical ER stress. Our previous findings indicate that when ER stress occurs in early cell cycle stages, before the ER stably enters the daughter cell and before initial ER tubule attachment to the polarisome at the bud tip, it leads to a block in ER inheritance into the daughter cell^25,31^. Once attached at the daughter cell bud tip, the direction of ER growth shifts laterally under the daughter cell’s plasma membrane. Ultimately, cells undergo cytokinesis to physically separate into two daughter cells^50^. In contrast, cells with larger bud sizes—indicating they are further along in the cell cycle—showed minimal impact on cER inheritance and continued to undergo cytokinesis even under canonical ER stress. However, in the second round of the cell cycle, cER inheritance was blocked at early stages of cell division^26,29^. Monitoring an asynchronous population of cells revealed that the percentage of cells experiencing cER inheritance block remained minimal as cells progressed to later stages of the cell cycle. Thus, among various cell cycle-impacted events, the cER inheritance block induced by *canonical ER stress* is a unique feature. It was surprising to find that Nsp4 expression-induced cER inheritance block occurs robustly even in class III yeast cells. Induction of the UPRE-GFP reporter via Nsp4 in yeast cells confirmed that ER stress is induced similarly to *canonical ER stress*.

The observed differences in the cER inheritance block suggest two possibilities: (1) ER stress induced by virally encoded proteins, such as Nsp4, may differ from canonical ER stress caused by endogenous proteins or cellular lipids. Previous studies have shown that viral proteins often display more disordered and loosely packed cores compared to host cell proteins, which may help them accommodate high mutation rates necessary for evading host defenses^51,52^. In addition, zoonotic transfers across species barriers may lead viral proteins to interact with host proteins from different species^53^. In contrast, host cell proteins are typically characterized by tightly packed core structures that provide thermodynamic stability. These structural differences may contribute to more significant ER stress and a more severe ER inheritance phenotype. Alternatively, (2) the ER inheritance block might be more sensitive to ER morphology, and the morphological changes induced by Nsp4 may result in a distinct ERSU mechanism. While ER morphological recovery in Vps4 knockout cells partially restores the ER inheritance block, future investigations into the ER inheritance behaviors in yeast cells with SARS-CoV-2 Nsp3 or Nsp6, or in combination, would be valuable.

We found that Nsp4 expression-induced ER morphological changes in yeast. In addition, while overall ER morphologies differ significantly between yeast and human cells, the basic nature of Nsp4-dependent ER morphological changes appears similar. Specifically, Nsp4 expression results in the formation of small ER foci in both yeast and human cells. Yeast ER contains more phosphatidylethanolamine, which is important for establishing membrane curvature facilitating lipid transfer. In contrast, the human ER membrane has a higher content of phosphatidylcholine compared to phosphatidylethanolamine. Thus, while similar ER associated curvatures may be generated by Nsp4, different ESCRT proteins may be involved for yeast or human ER morphological changes. Future work will require to determine if ESCRT protein is involved in Nsp4-induced ER structural changes and ultimately in DMV formation. A recent report on *in vitro* reconstitution, demonstrating the dynamic assembly and disassembly of specific ESCRT protein subunits, identified at least three major intermediate stages for achieving ESCRT protein functions^36,54,55^: (1) association of a subset of ESCRT subunit affinities, (2) the specific affinity between subunits facilitating subsequent recruitment, and (3) the role of Vps4 in driving unidirectional progression of ER membrane changes through ATP hydrolysis. Based on the proposed ESCRT-III polymerization model, the affinities between Vps23, Vps24, and Snf7—forming a core ESCRT-III protein complex— may drive ER membrane changes, leading to further changes mediated by ATP-dependent Vps4^56^. Further work is needed to identify if and which specific ERSCRT proteins are involved in the Nsp4- induced ER structural changes in human cells. In addition, future work is needed to identify the components involved in the Nsp4-induced ERSU pathway, especially since knockout of Slt2 did not affect the ER inheritance block, unlike the role of SLT2 in canonical ER stress^26^. Furthermore, the reasons for these differences in ER inheritance block behaviors and their underlying molecular mechanisms will wait for further investigation. The experimental system we have developed is well-suited to reveal previously uncharacterized differences in ER responses to proteins and lipids from host pathogens.

Recent studies have linked SARS-CoV-2 or its viral family members to the development of acute respiratory distress syndrome (ARDS), which induces the UPR pathway^57,58^, including ATF6. Our study suggests that SARS-CoV-2 Nsp4 has unique features that induce ER stress and UPR activation. While the detailed molecular mechanisms of ATF6 activation by SARS-CoV-2 will require future studies, drugs that modulate ATF6 have shown promise for treating ARDS. Thus, the experimental system we have developed is poised to uncover previously uncharacterized differences in ER responses to proteins and lipids from host pathogens, including SARS-CoV-2.

## Materials and Methods

### Strains and expression of SARS-CoV-2 Nsp4 in yeast cells

Details of the *S. cerevisiae* strains used in this study, generated using primers described in Table 2, are provided in Table 1. Primers shown in Table 2 were used to generate . Yeast cells expressing SARS-CoV-2 Nsp4 were grown in synthetic complete -URA media containing 2% raffinose. Expression of Nsp4 was induced upon switching to growth media containing 2% Gal (+ Gal) for 3 hr. All experiments involving expression of Nsp4 were performed using the same approach. The Nsp4 plasmid (pGBW-m4046479) containing a URA auxotroph was obtained from AddGene and transformed into all strains in Table 3. HeLa cells stably express Sec61β-GFP ER reporter were generated from HeLa cells (obtained from ATCC, CCL-2). U2OS cells stably express mEmerald- Sec61β were obtained from Dr. Craig Blackstone (Harvard MGH, Boston) and the source of the cell line carrying this reporter was described in their publication^59^.

**Table 1:** Yeast Strains.

| Strain Name | Genotype | Reference |
| --- | --- | --- |
| BY4741 | *MATa his3Δ1, leu2Δ0, met15Δ0, ura3Δ0* | Thermo Fisher Scientific |
| BY7092 | *MATa can1Δ::STE2pr-Sp_his5, lyp1Δ, his3Δ1, leu2Δ0, ura3Δ0, met15Δ0* | Boone Lab |
| MNY2661 | *BY4741 MATa his3Δ1, leu2Δ0, met15Δ0, ura3Δ0:: PHO88-GFP:HIS3* | Niwa lab |
| MNY2817 | *BY7092 MATa can1Δ::STE2pr-Sp_his5, lyp1Δ, his3Δ1, leu2Δ0, ura3Δ0, met15Δ0:: PHO88-GFP:HIS3, slt2Δ::Nat* | Niwa lab |
| MNY3167 | *BY4741 MATa his3Δ1, leu2Δ0, met15Δ0, ura3Δ0::PHO88-GFP:HIS3, ire1Δ::KanMX* | Niwa lab |
| *vps4Δ* | *BY4741 MATa his3Δ1, leu2Δ0, met15Δ0, ura3Δ0:: PHO88-GFP:HIS3, vps4Δ::KanMX* | Thermo Fisher Scientific |
| *vps24Δ* | *BY4741 MATa his3Δ1, leu2Δ0, met15Δ0, ura3Δ0:: PHO88-GFP:HIS3, vps24Δ::KanMX* | Thermo Fisher Scientific |
| *vps23Δ* | *BY4741 MATa his3Δ1, leu2Δ0, met15Δ0, ura3Δ0:: PHO88-GFP:HIS3, vps23Δ::KanMX* | Thermo Fisher Scientific |
| *snf7Δ* | *BY4741 MATa his3Δ1, leu2Δ0, met15Δ0, ura3Δ0:: PHO88-GFP:HIS3, snf7Δ::KanMX* | Thermo Fisher Scientific |
| *ubx2Δ* | *BY4741 MATa his3Δ1, leu2Δ0, met15Δ0, ura3Δ0:: PHO88-GFP:HIS3, ubx2Δ::KanMX* | Thermo Fisher Scientific |
| *mga2Δ* | *BY4741 MATa his3Δ1, leu2Δ0, met15Δ0, ura3Δ0:: PHO88-GFP:HIS3, mga2Δ::KanMX* | Thermo Fisher Scientific |

**Table 2:**
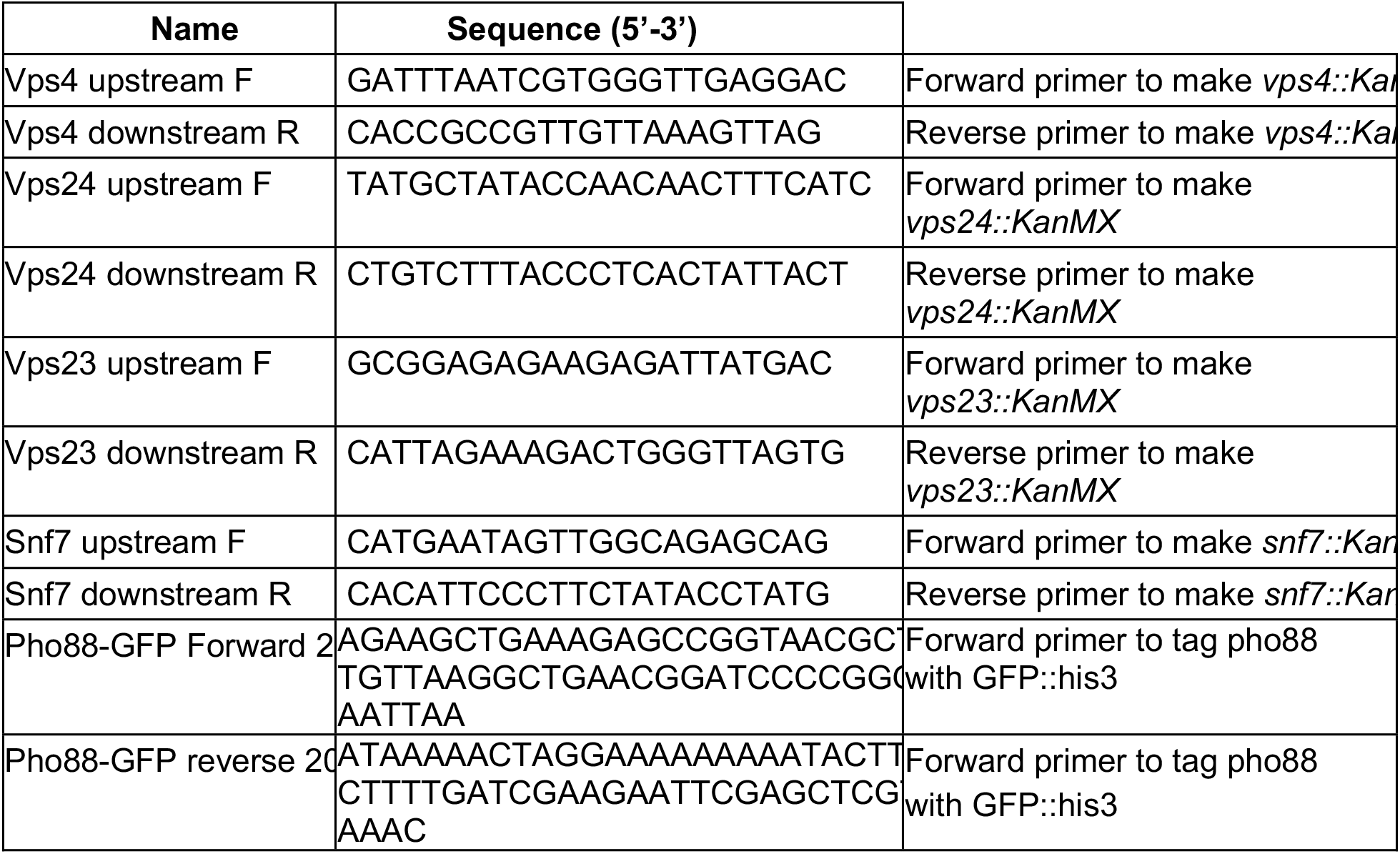
Primers.

**Table 3:** Plasmids.

| Plasmid Name | plasmid |  |
| --- | --- | --- |
| For tagging yeast gene with GFP at genomic locus | pFA6a-GFP (S65T)-His3MX6 |  |
| For Nsp4 ura | pGBW-m4046479 | Addgene |

### Total cell extract and Western blot

Yeast cells were harvested by centrifugation and resuspended with lysis buffer (20 mM HEPES, pH 7.4, 150 mM NaCl, 1% Triton-X, 10% glycerol, 1 mM EDTA, 100 mM NaF, 1mM PMSF) containing protease inhibitor cocktail (Gold Bio) and bashed with acid-washed beads for 10 min at 4°C. Protein concentration was determined using a Pierce™ BSA Protein Assay (Thermo-Fisher). A total of 20 mg of total extract was loaded on 10% SDS-PAGE and blotted on nitrocellulose membrane using a wet transfer at 100V for 60 min. Membrane was Ponceau stained for assessment of protein levels in each lane. The primary antibody used was anti-Flag M.2 antibody (Millipore-Sigma) at a 1:1,000 dilution for a 2 hr incubation at room temperature. The secondary antibody was HRP-conjugated goat anti-mouse antibody (Bio-Rad) used at a 1:5,000 dilution and incubated for 1 hr at room temperature. Membranes were developed with Super Signal West Pico PLUS Chemiluminescent Substrate (Thermo-Fisher) and imaged using a Chemi Doc XRS+ (Bio-Rad).

### FACS analyses of Nsp4-RFP

At 18 hr post transfection, U2OS cells with Nsp4-RFP were treated with 0.05% Trypsin-EDTA to detach cells from wells. Cells were collected via centrifugation and washed in 2 mL of cold 3% FBS-PBS solution. After washing, cells were resuspended in a final volume of 300 mL of 3% FBS-PBS solution and passed through 40 mM cell strainers into FACS tubes and kept on ice until analysis.

The levels of Nsp4-mCherry in U2OS cells were measured by flow cytometry using a CytoFlex S flow cytometer (Beckman Coulter) and analyzed using FlowJo™ software (BD Life Sciences).

### ER stress induction by tunicamycin

ER stress was induced upon addition of Tm (Calbiochem) at a final concentration of 1 μg/mL at 30°C for 2 hr to analyze ER morphology and morphological changes. Tm (-) cells were treated with DMSO. To visualize the nucleus and bud scars or CRMs, we used DAPI and 0.1 μg/ml Calcofluor White, CFW, (Sigma), respectively. For staining with CFW, cells were washed once with 1X phosphate buffered saline (PBS), incubated with CFW for 2 min in the dark, and then washed once more with 1X PBS. Cells were imaged live immediately after staining.

### Fluorescence microscopy

All live cells were visualized using an Axiovert 200M Carl Zeiss Micro-Imaging microscope with a 100x 1.3 NA objective. Images were captured using a monochrome digital camera (Axiocam; Carl Zeiss Micro Imaging, Inc.) and analyzed using Axiovision software (Carl Zeiss Micro Imaging, Inc.). Images were acquired with a monochrome digital camera with a z-stack projection of 0.11-µm intervals (Axiocam; Carl Zeiss Micro Imaging) and analyzed after constrained iterative deconvolution using Zen software (Carl Zeiss Micro imaging).

### Quantification and statistical analysis

ImageJ software (National Institute of Health) was used for all quantification. A minimum of 150 cells were measured for each experiment and each experiment was repeated three times, unless otherwise stated. Progression through the cell cycle was determined by the bud index (ratio of daughter and mother cell diameters) and is classified as follows: Class I cells (bud index of 0–0.3), Class II cells (bud index of 0.3–1 with no pnER divided into the daughter), and Class III cells (bud index of 0.3–1 with pnER divided into the daughter), as described previously^26^. Cell diameters were measured in ImageJ. Daughter buds were scored for the inheritance of cER if GFP signal lined at least 80% of the bud cortex. Nsp4-induced changes on the ER structure were shown by # of cells with altered morphology. Quantitative data are expressed as mean ± SD. Student’s two-tailed T test was used to generate *p* values.

### SARS-CoV-2 expression in HeLa cells and siRNA treatment

HeLa cells stably expressing Sec61b-GFP were placed in 6-well plates containing poly-L-lysine–treated cover slips and grown to 70% confluency. Cells were transfected with 2,400 ng of plasmid using the Effectene Transfection Reagent (Qiagen), according to the manufacturer’s instructions. U2OS or HeLa cells were transfected with 2,400 ng of Nsp4-RFP or Nsp4 plasmid using the Effectene Transfection Reagent (Qiagen) according to the manufacturer’s instructions. At 18 hr post-plasmid transfection, cells were treated with DAPI and fixed with 4% paraformaldehyde (PFA; Sigma P6148) for imaging analysis.

### Immunofluorescence

For immunofluorescence microscopy, cells were treated and fixed with 4% PFA (Sigma P6148) for 5 min, permeabilized with 0.2% NP-40 in PBS for 5 min, and blocked in 5% BSA in PBS for at least 1 h. The primary anti-GM130, anti-GPP130, anti-Rtn4 antibodies, followed by secondary antibody (Alexa Fluor 568, Life Technologies), were used for visualization of the Golgi. For visualization of the nucleus, 1 mg/ml DAPI (Pierce) was added to the mounting media (Aqua-Mount; Lerner Labora-tories). Coverslips were mounted onto slides and the cells were imaged with an Observer Z1 microscope (Carl Zeiss MicroImaging).with either a 100X or a 633 1.4 NA objective.

## Data Availability

The datasets used and/or analysed during the current study available from the corresponding author on reasonable request.

